# Investigation of Powassan virus lineage II pathogenesis and neurotropism in mice

**DOI:** 10.1101/2025.11.07.687151

**Authors:** Samantha J. Courtney, Emily N. Gallichotte, Emma Nilsson, Chasity E. Trammell, Kate X. Kimball, Anna C. Fagre, Allison Vilander, Anna K. Överby, Gregory D. Ebel

**Affiliations:** Department of Microbiology, Immunology and Pathology, Colorado State University, Fort Collins, CO 80523, USA; Department of Clinical Microbiology, Umeå University, 90185, Umeå, Sweden; Laboratory for molecular infection medicine Sweden (MIMS), Umeå University, 90187, Umeå, Sweden

## Abstract

Powassan virus (POWV) is an emerging tick-borne flavivirus that causes disease in humans. POWV has considerable genetic and phenotypic diversity, including highly variable replication *in vitro* and pathogenesis in mice. This study sought to define the extent of variability in pathogenesis within POWV lineage II in mice and investigate possible viral determinants. Relative to other strains, two New York-derived isolates, NY.19.12 and NY.19.32, caused earlier clinical signs and earlier detection of viral RNA (vRNA) in the spleen and brain compared to mice infected with virus derived from a lineage II infectious clone (WI.97.ic). Sequencing revealed these strains share three amino acid substitutions in envelope, NS1, and NS5 compared to other lineage II strains, which were engineered into a mutant infectious clone. At early time points post-infection, clinical signs, vRNA detection in the cerebellum, and viral distribution in the brain were similar between NY.19.12 and the mutant clone, suggesting these mutations may play a role in disease progression and early neuroinvasion. However, NY.19.12 vRNA was detected in the spleen at significantly higher rates compared to both WI.97.ic and the mutant clone, indicating factors other than these mutations are responsible for increased spleen infection. Importantly, this study highlights the complexity of POWV pathogenesis and suggests that POWV lineage II strains have varying disease phenotypes likely driven by multiple genetic differences.

**Importance:** Tick-borne flaviviruses exhibit considerable genetic and phenotypic diversity in nature, influencing their transmission and pathogenesis. Defining the mechanisms of pathogenesis requires understanding how inter-strain variation translates to phenotypic differences in viral dissemination and neuroinvasion. This study demonstrates that even closely related Powassan virus (POWV) lineage II strains display distinct disease phenotypes that are only partially attributable to nonsynonymous consensus mutations within the viral coding sequence. By highlighting the complexity of POWV infection dynamics in mice, these findings provide valuable insights into how POWV lineage II diversity may shape disease progression and severity in humans.

## Introduction

Powassan virus (POWV, Flaviviridae, *Orthoflavivirus*) is a tick-borne flavivirus that causes severe neuroinvasive disease in humans.^1^ POWV is most frequently detected in the upper Midwest and Northeastern United States, Canada and the Russian Far East.^2^ POWV is classified into two distinct lineages. Lineage I appears to mainly be associated with *Ixodes cookei* and *I. marxi*. Lineage II is mostly associated with *I. scapularis*, which frequently feed on humans.^3,4^ POWV transmission in nature is highly focal, with genetically similar strains deriving from the same geographic region.^5^ However, there is limited understanding of strain-specific impacts on virus transmission by ticks and disease outcomes in humans.

Following the bite of an infected tick, symptom onset in humans often begins with flu-like illness including fever and vomiting. In severe cases, symptoms progress to encephalitis characterized by headaches, confusion, loss of coordination, and speech difficulties.^2,6^ Ten percent of clinically apparent cases are fatal and neurologic sequelae persist in up to 50% of symptomatic cases.^7^ There is an urgent need to better understand the pathogenesis of this emerging tick-borne virus because the number of reported human cases has markedly increased over the last 20 years.^8^

Mice are widely used as an experimental model for POWV infection and pathogenesis, where peripheral inoculation routes such as subcutaneous, intradermal, or footpad injections are designed to mimic natural tick-bite transmission. The virus replicates to high levels in the central nervous system (CNS), leading to significant neuropathology and a high mortality rate in C57BL/6 mice.^9^ In addition, there is evidence of virus accumulating in peripheral tissues such as the lymph nodes and spleen.^10^ However, differences in the pathogenesis and tropism of genetically diverse POWV lineage II strains have not been extensively studied in mice. Additionally, there is limited research on specific viral determinants of POWV neuroinvasion and disease.

We therefore sought to determine the degree of variation in pathogenesis among POWV lineage II virus strains using C57BL/6 mice. Several isolates from different geographical regions and a Midwest lineage II infectious clone-derived virus were selected, and morbidity, mortality, and tissue tropism were assessed. Significant variation in pathogenesis was observed, with two strains from New York demonstrating enhanced infection of the spleen and brain. These strains shared three unique amino acid substitutions in envelope (E), non-structural protein 1 (NS1), and NS5 proteins. When the mutations were engineered into an infectious clone, they recapitulated the early neuroinvasion phenotype but not spleen tropism. Overall, these results establish that POWV pathogenic potential may be highly strain-dependent and demonstrate the complexity of genotypic differences impacting variation within POWV lineage II disease phenotypes in mice.

## Results

### Selected POWV lineage II strains

POWV lineage II strains group phylogenetically based on geographic origin (**Figure 1A**).^5^ Virus strains for pathogenesis studies were selected to represent the broad geographic range and genetic diversity within lineage II (**Figure 1B**, **Table 1**).

**Figure 1.**
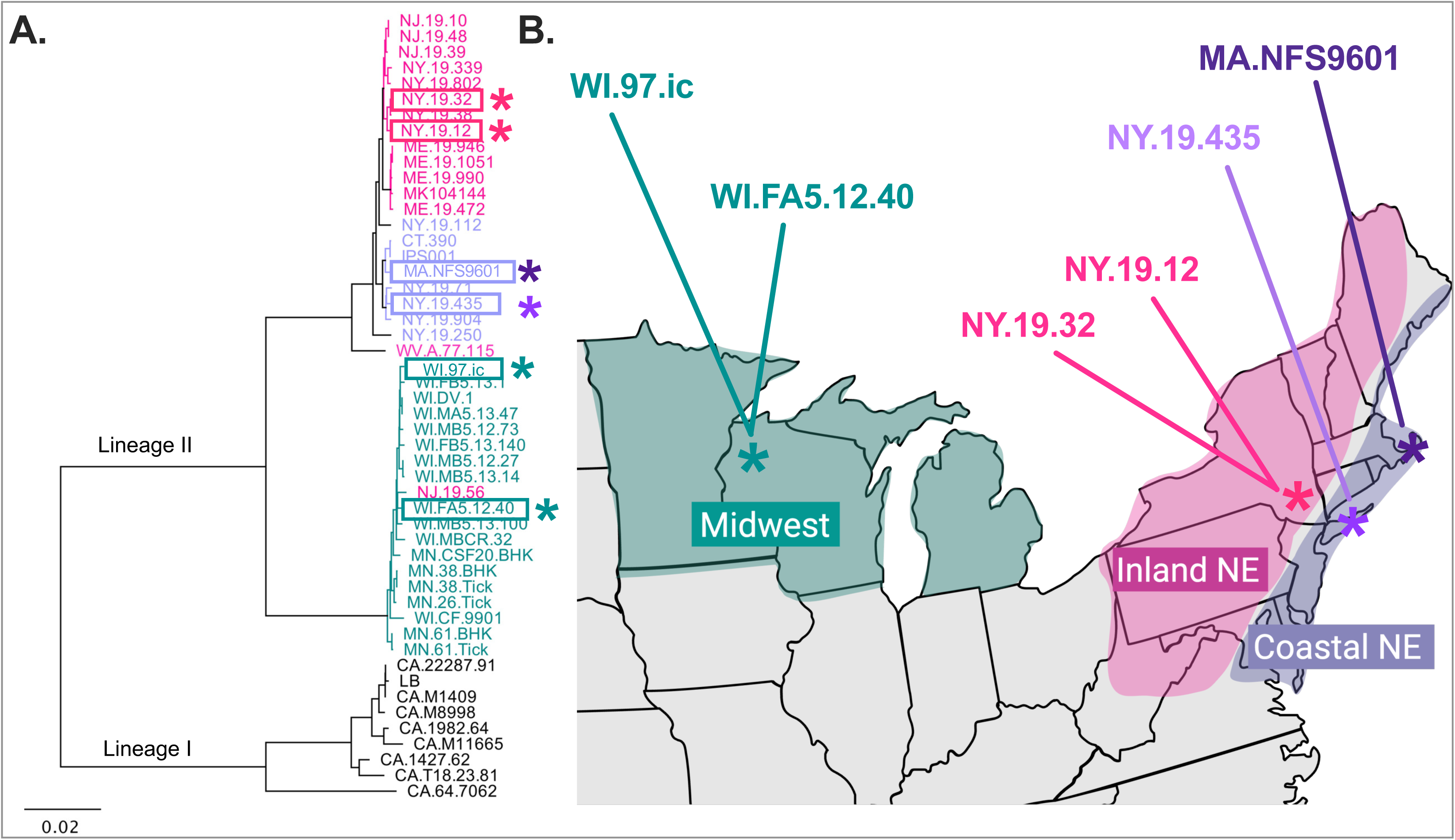
POWV lineage II strains selected based on geography. **A)** A rooted neighbor-joining phylogenetic tree of full genome amino acid sequences of POWV lineage I and II sequences generated in Geneious Prime ® 2023.0.1 (black = lineage I; purple = lineage II, coastal Northeast; pink = lineage II, inland Northeast; teal = lineage II, Midwest). Viruses used in subsequent studies are boxed. **B)** United States map showing location where each lineage II POWV strain was originally collected, color-coded as described in A (asterisks correspond to selected strains in the phylogenetic tree).

**Table 1.** Characteristics of POWV lineage II strains used.

| Strain Designation | Source Location | Year of isolation | Species Origin | Passage History | Accession number |
| --- | --- | --- | --- | --- | --- |
| MA.NFS9601 | Nantucket, MA | 1996 | *I. scapularis* | SM1B2 | HM440559 |
| NY.19.435 | Connetquot, NY | 2019 | *I. scapularis* | B2 | OP823469 |
| WI.FA5.12.40 | Spooner, WI | 2008 | *I. scapularis* | B2 | OP823475 |
| WI.97.ic | Spooner, WI | 1997 | *I. scapularis* | B3 | OP823482* |
| NY.19.32 | Saratoga Spa, NY | 2019 | *I. scapularis* | B2 | OP823462 |
| NY.19.12 | Saratoga Spa, NY | 2019 | *I. scapularis* | B2 | OP823461 |

\*WI.97.ic is a clone-derived virus from this GenBank accession number sequence. SM = suckling mice, B = BHK cells

### POWV morbidity and mortality in mice

Differences in morbidity and mortality in C57BL/6 mice using genetically diverse virus strains were assessed. Survival of mice inoculated subcutaneously (S.C.) with MA.NFS9601, NY.19.435, WI.FA5.12.40, or WI.97.ic ranged from 25-75% over 14 days (**Figure 2A**). Conversely, by seven days post-infection, 75% and 100% of NY.19.32 and NY.19.12 infected mice, respectively, had succumbed to disease (**Figure 2A**). Survival was significantly lower in NY.19.12 infected mice compared to WI.97.ic mice (p<0.0001). By day three post-infection, NY.19.12 and NY.19.32 infected mice displayed mild clinical signs, and by day seven post-infection, 78% required euthanasia, compared to only 22% of mice infected with other POWV strains (**Figure 2B**). By day six post-infection, all NY.19.12 infected mice were exhibiting signs of forelimb weakness (**Figure 2C**). Additionally, most NY.19.12 and NY.19.32 infected mice rapidly lost weight within seven days of infection (**Figure 2D**), whereas weight loss for other groups varied, with many mice recovering following initial weight loss (**Figure 2D**).

**Figure 2.**
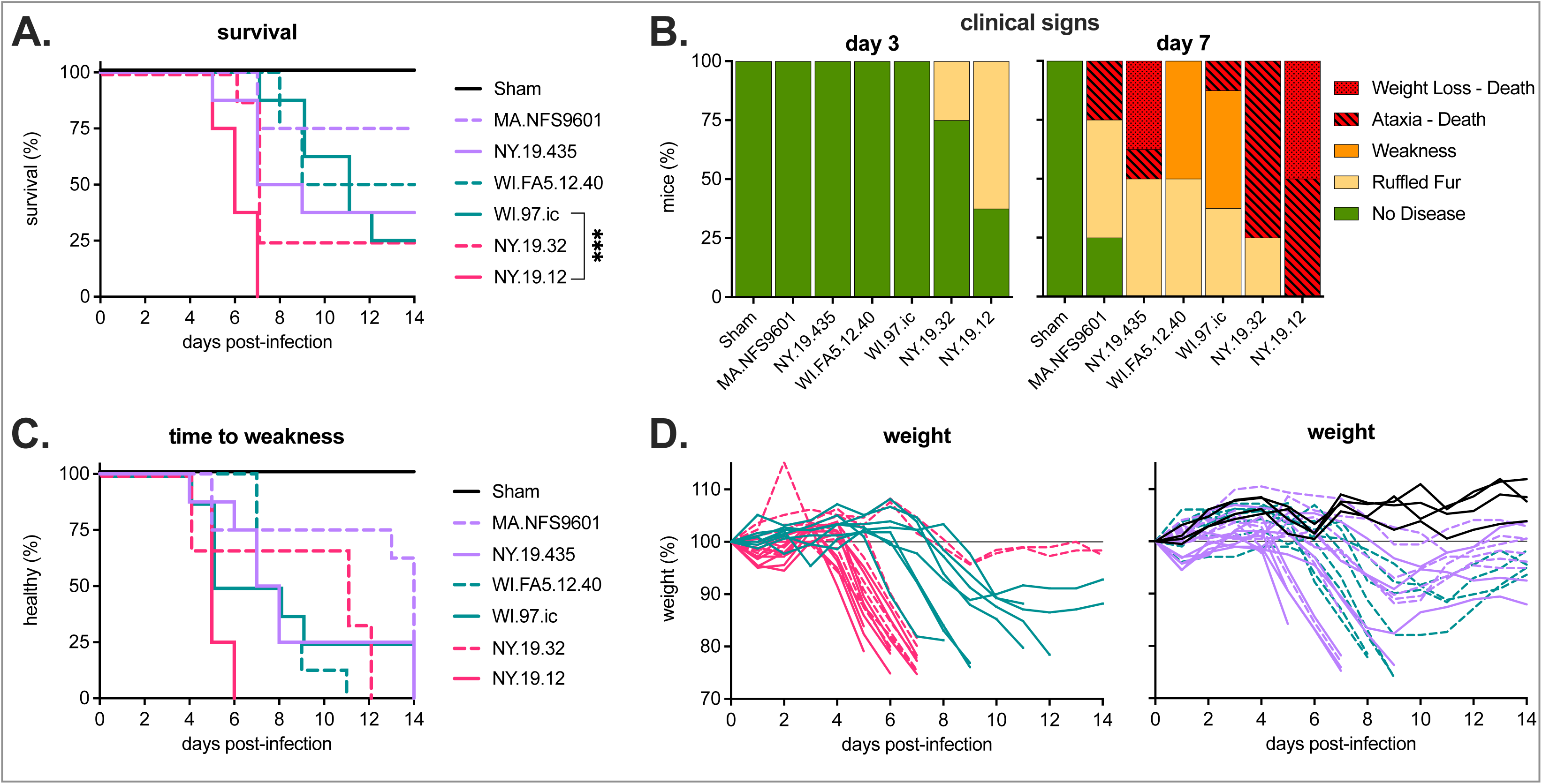
NY.19.12 and NY.19.32 cause increased morbidity and mortality in mice. Mice were S.C. inoculated with 10^3^ PFU POWV (MA.NFS9601, NY.19.435, WI.FA5.12.40, WI.97.ic, NY.19.32, NY.19.12) or sham and monitored daily for 14 days (n = 8 per strain). **A)** Kaplan-Meier survival curves. Survival of each virus was compared to WI.97.ic, with log-rank (Mantel-Cox) test, *** p<0.0001. **B)** Clinical signs on days three and seven post-infection. **C)** The amount of time for mice to exhibit forelimb weakness. No comparisons between each virus and WI.97.ic were significant using log-rank (Mantel-Cox) test, p>0.05. **D)** Percent weight change from starting weight of NY.19.12, NY.19.32, and WI.97.ic infected mice, and MA.NFS9601, NY.19.435, WI.FA5.12.40, and sham infected mice.

### Viral tissue tropism in mice

To determine POWV infection kinetics and tissue tropism for NY.19 isolates and WI.97.ic, C57BL/6 mice were S.C. inoculated and sampled over ten days. Viral loads were assessed in serum, spleen and cortical tissue of the brain. For mice sacrificed between one to six days post-infection, there was no weight loss (**Figure 3A**). At seven days post-infection, only mice infected with NY.19.12 and NY.19.32 had lost weight (**Figure 3A**) and 15% met endpoint criteria for this study (ataxia or 20% weight loss), compared to WI.97.ic infected mice, which only displayed mild to moderate clinical signs (**Figure 3B**). Viremia was detected more frequently in NY.19.12 and NY.19.32 infected mice compared to WI.97.ic, most notably between one to four days post-infection, and significantly more frequently across all timepoints combined (p<0.05) (**Figure 3C**). In addition, NY.19.12 viral RNA (vRNA) was detected significantly more frequently than WI.97.ic in spleens throughout the course of infection (p<0.01) (**Figure 3D**) and in the cortex at earlier time points (days five and six) compared to WI.97.ic (day eight) (**Figure 3E**). Despite lower rates of viremia and later neuroinvasion in WI.97.ic infected mice, levels of vRNA in serum and cortex were similar to NY.19.12 and NY.19.32 (**Figure 3F/H**). In the spleen, however, average vRNA was significantly higher in NY.19.12 and NY.19.32 infected mice compared to WI.97.ic (p<0.05) (**Figure 3G**). Given the genotypic and phenotypic similarities between NY.19.12 and NY.19.32, further analyses proceeded with only NY.19.12. The right side of the brain from mice that succumbed to disease by day ten were characterized via histopathology. Overall, lesions in brains from both WI.97.ic and NY.19.12 were similar. In the cerebrum of all infected brains, there was lymphohistiocytic meningitis to meningoencephalitis with gliosis and neuronal degeneration and necrosis (**Figure 4)**.

**Figure 3.**
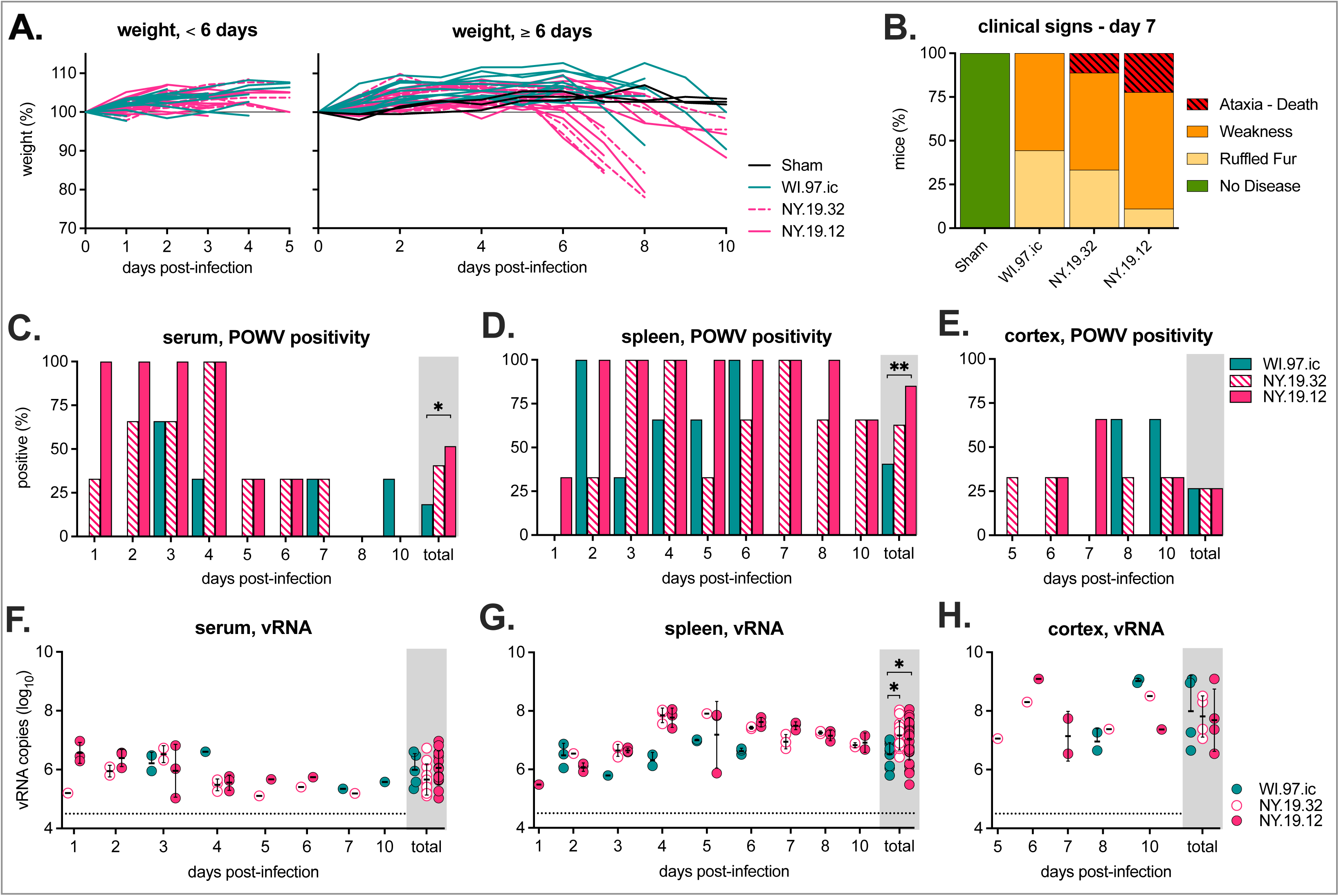
NY.19.12 and NY.19.32 mice succumb to disease earlier and have higher rates of tissue infection. Mice were S.C. inoculated with 10^3^ PFU POWV (NY.19.12, NY.19.32, WI.97.ic) or sham, sacrificed, and sampled over a ten-day course of infection (n = 27 per infection cohort, 3 mice sacrificed per time point). **A)** Percent weight change from starting weight for mice sacrificed days one through five post-infection (n = 15) and those sacrificed post-day six (n = 12). **B)** Clinical signs seven days post-infection (n = 9). Positive POWV (%) **C)** serum, **D)** spleen and **E)** cortex per day and overall (Fisher’s exact test, * p<0.05, ** p<0.01). vRNA copies per **F)** milliliter of serum, and gram of **G)** spleen, and **H)** cortex per day post-infection (mean ± standard deviation). The dotted line shows the limit of detection. For each sample type, one-way ANOVA Kruskal Wallis test on total samples (* p<0.05).

**Figure 4.**
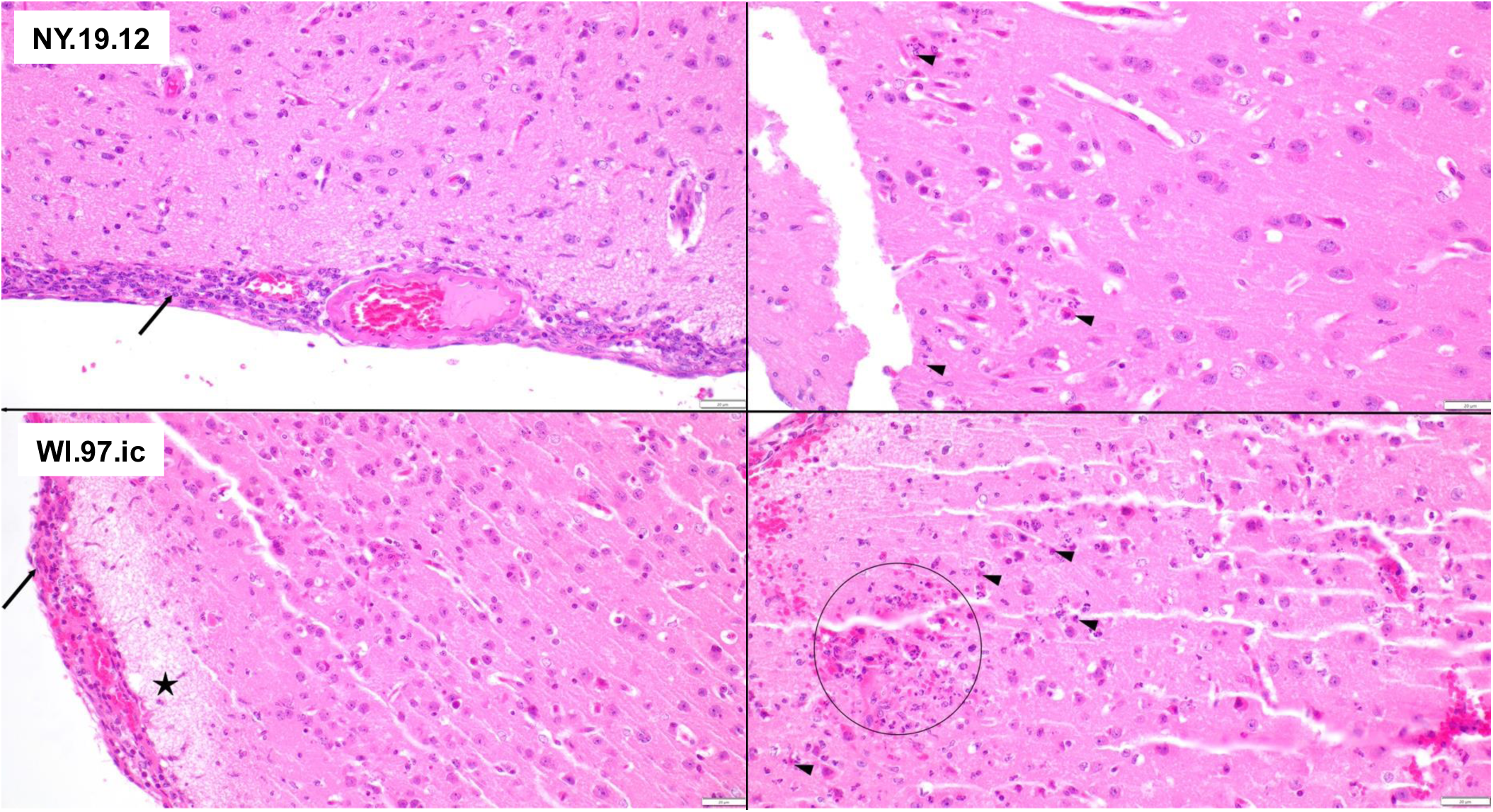
Infected mouse brains showed similar severity and distribution of inflammation. Representative histopathology of NY.19.12 (top) and WI.97.ic (bottom) infected mouse brains. There is meningitis (arrows), superficial cerebral edema (star), neuronal degeneration and necrosis (arrow heads), and neutrophilic inflammation with hemorrhage (circle). Scale bars = 20 µm. H&E.

### Viral RNA in distinct brain regions

POWV lineage I and other tick-borne flaviviruses frequently infect distinct brain regions.^10,11^ However, this has not been well characterized for POWV lineage II.^12,13^ Therefore, we assessed potential differences in viral distribution in the brain at disease onset (i.e. six days post-infection). We measured vRNA in four distinct brain regions (olfactory bulbs, cortex, cerebellum, and brainstem) of mice infected with NY.19.12 and WI.97.ic (**Figure 5A**). No mice met criteria for euthanasia at six days post-infection (**Figure 5B/C**). However, at this timepoint, 100% of NY.19.12 infected mice exhibited clear neurological signs (i.e., forelimb weakness), compared to 33% of WI.97.ic infected mice (**Figure 5D**), and NY.19.12 infected mice displayed weakness earlier than those infected with WI.97.ic (**Figure 5E)**. In all NY.19.12 and WI.97.ic infected mice, vRNA was detected at 6 days post-infection in the brainstem, cortex, and olfactory bulbs (**Figure 5F**), but NY.19.12 had higher rates of virus detection in the cerebellum and the spleen (**Figure 5F**). In addition, NY.19.12 had higher levels of vRNA in the spleen but differences were not significant (**Figure 5G**).

**Figure 5.**
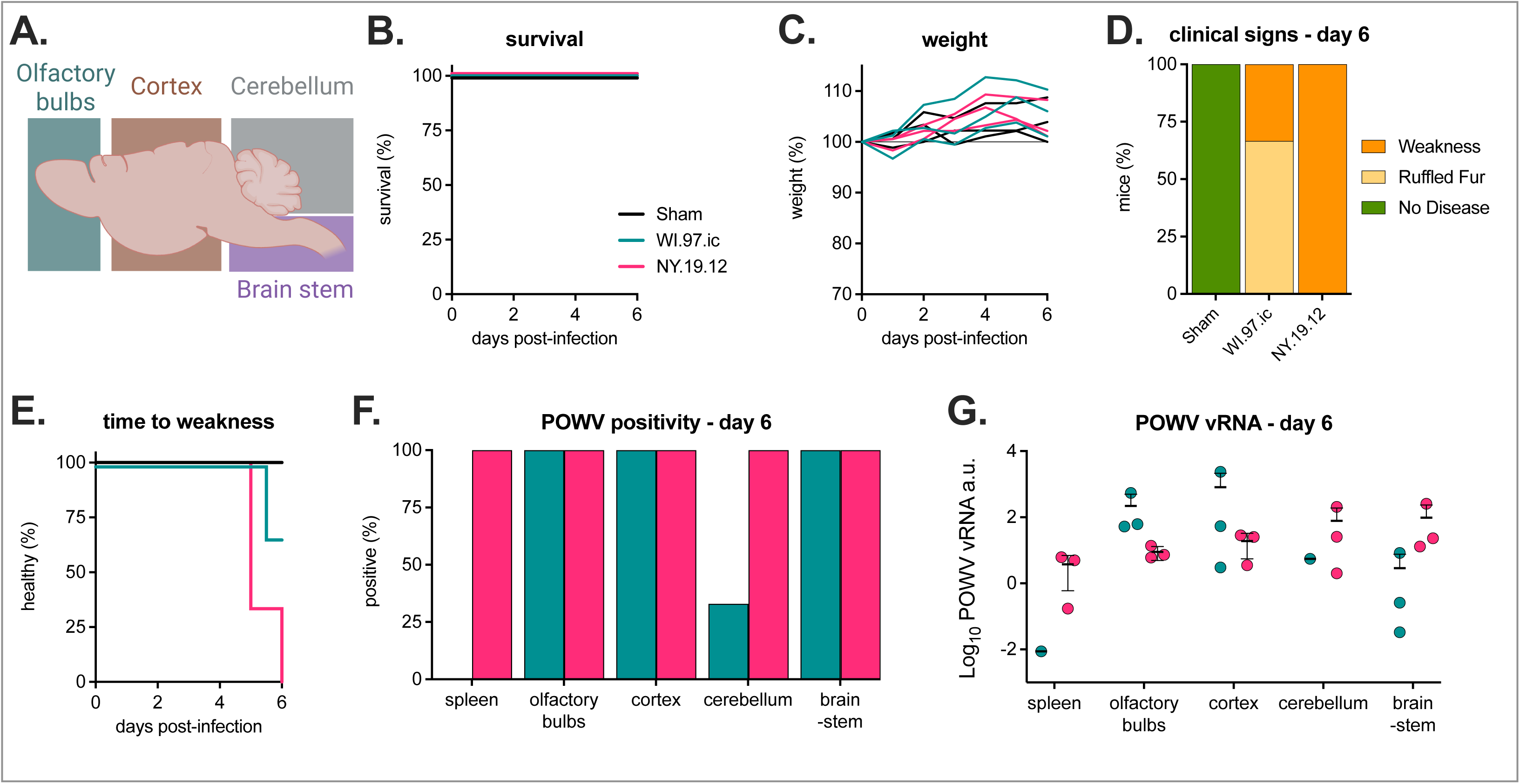
NY.19.12 infected mice display early neurological signs and vRNA appears more frequently in cerebellum and spleen. Mice were infected with POWV (NY.19.12, WI.97.ic) or sham and sacrificed at six days post-infection (n = 3 per cohort). **A)** Brain regions that were harvested (olfactory bulbs, cortex, cerebellum, and brainstem). **B)** Kaplan-Meier survival curve. **C)** Percent weight change from starting weight. **D)** Clinical signs six days post-infection. **E)** The amount of time for mice to exhibit forelimb weakness. Comparison of WI.97.ic and NY.19.12 was not significant using log-rank (Mantel-Cox) test, p>0.05. **F)** Positive POWV (%) samples from spleen, olfactory bulbs, cortex, cerebellum, and brainstem (no significant differences by Fisher’s exact test). **G)** Log_10_ POWV vRNA normalized to GAPDH intracellular levels from spleen, olfactory bulbs, cortex, cerebellum, and brainstem. a.u. = arbitrary units. No significant differences within sample type by Mann Whitney t-test (p>0.05).

To further characterize neurotropism, mice were S.C. inoculated with NY.19.12 and sacrificed at day six post-infection, where brains were processed for an immunofluorescence assay (IFA) targeting POWV, GFAP (astrocytes), and NeuN (neurons). There was minimal colocalization of NY.19.12 with astrocytes (**Figure 6A/B**). However, colocalization of NY.19.12 with neuronal cells was observed, indicating that POWV specifically infects neurons (**Figure 6C/D)**.

**Figure 6.**
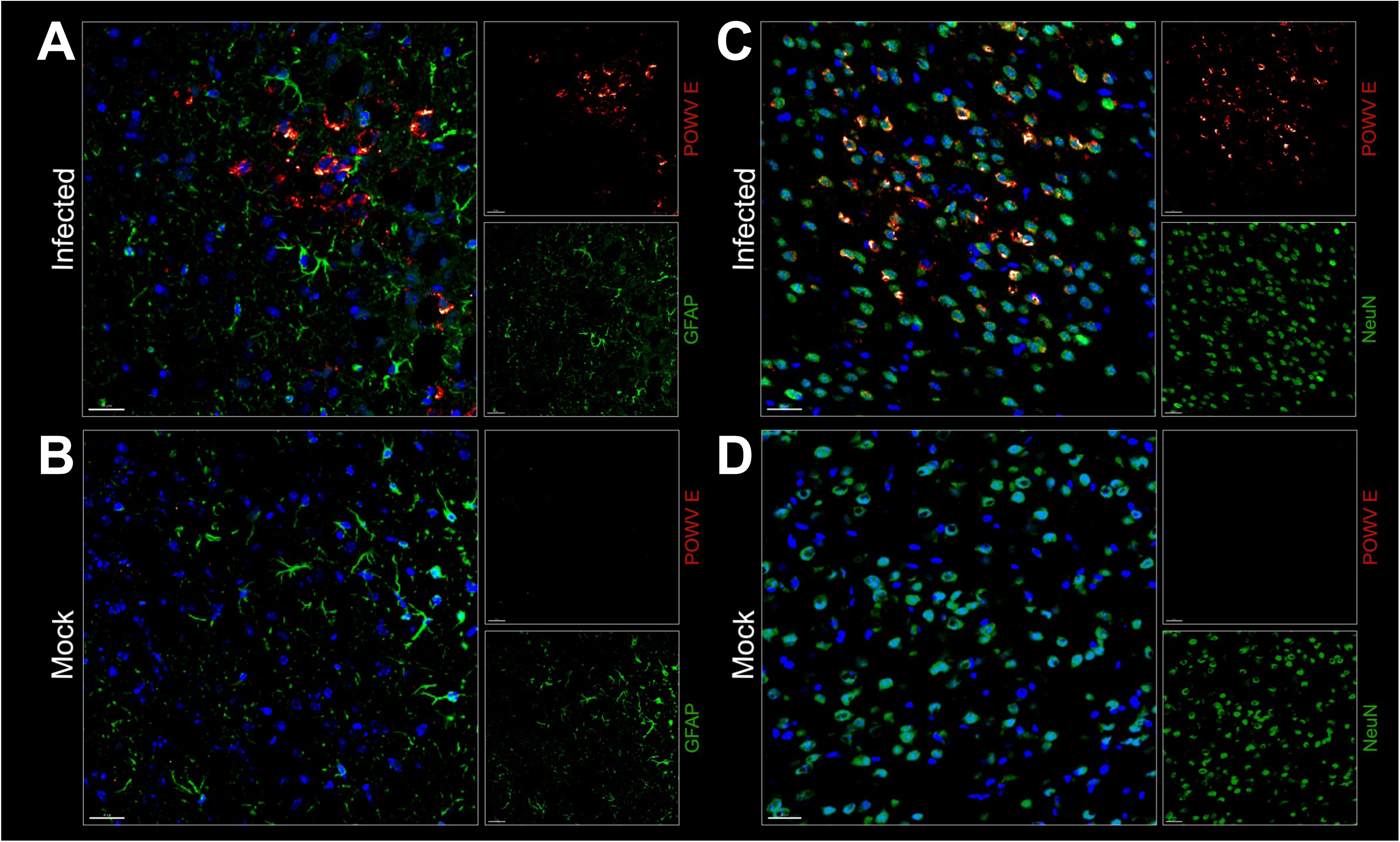
NY.19.12 specifically infects neurons with minimal astrocyte recruitment. Mice were infected with NY.19.12 and sacrificed at six days post-infection, where infected brains were processed for IFA. Representative axial images show: **A)** an NY.19.12 infected mouse brain stained for GFAP (astrocytes) in green and POWV E in red; **B)** an uninfected mock brain stained for GFAP and POWV E**; C)** an NY.19.12 infected mouse brain stained for neurons (NeuN) in green and POWV E in red and **D)** an uninfected mock brain stained for NeuN and POWV E.

### Genetic differences between strains

To identify genetic differences between virus strains, the amino acid sequences of all POWV strains used in the morbidity and mortality study were aligned (see **Table 1**). NY.19.12 and NY.19.32 share three unique amino acid substitutions in E, NS1 and NS5 (**Figure 7A**). These residues (E 205, NS1 178, NS5 649) were then examined in lineage I and lineage II POWV strains to define their prevalence among POWV in nature. The E and NS1 mutations were only detected in lineage II viruses, predominantly those from the inland Northeast (**Figure 7B**). While the NS5 mutation was detected in those viruses, it was also detected in every lineage I strain examined (n=50). To determine whether these mutations were maintained post-neuroinvasion, POWV was sequenced from cortical tissue of WI.97.ic, NY.19.32, and NY.19.12 infected mice (**Figure 3E/H**). All three mutations were maintained post-neuroinvasion, and WI.97.ic did not acquire them (**Table 2**). To investigate the role of these mutations in pathogenesis in mice, they were engineered into a WI.97.ic infectious clone backbone, designated WI.97.ic^NY.19.12^ (**Figure 7C**).

**Figure 7.**
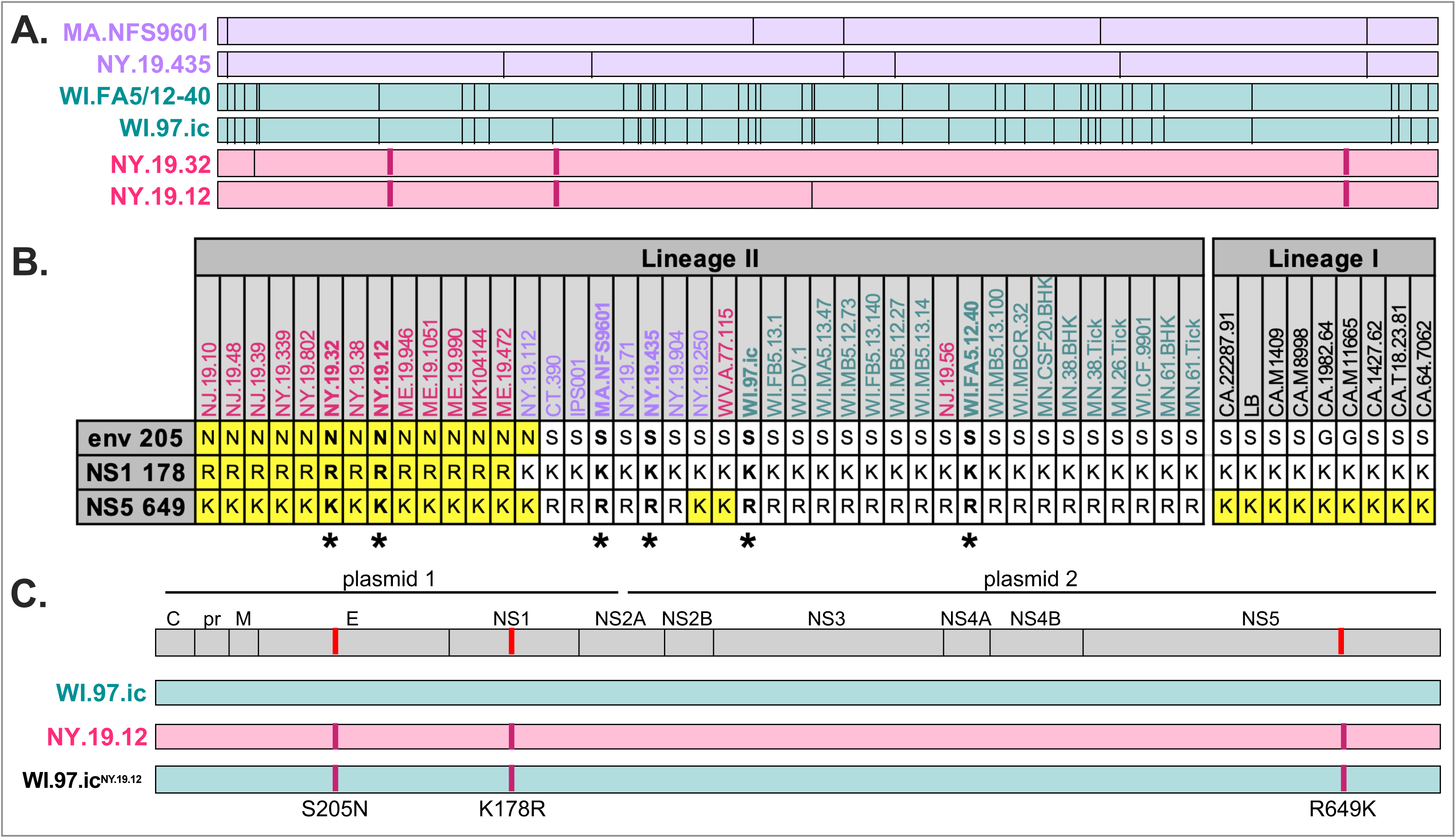
NY.19.32 and NY.19.12 share three amino acid mutations. **A)** Amino acid alignment of MA.NFS9601, NY.19.435, WI.FA5.12.40, WI.97.ic, NY.19.32, and NY.19.12. Amino acid differences between strains shown as lines. Three amino acids shared by NY.19.32 and NY.19.12 shown in dark pink. **B)** Amino acid residues at E-205, NS1-178, and NS5-649 among POWV lineage I and lineage II strains. Strains are ordered and colored based on the phylogenetic tree (see Figure 1A). Highlighted residues indicate the presence of the mutation (E-S205N, NS1-K178R, NS5-R649K). Strains labeled with * are viruses used in the morbidity and mortality study (see **Table 1**). **C)** E-S205N, NS1-K178R, NS5-R649K mutations were engineered into the WI.97.ic two-plasmid backbone, producing a mutant infectious clone (WI.97.ic^NY.19.12^).

**Table 2.** Sequence of POWV in cortical tissue.

|  |  | E |  | NS1 |  | NS5 |  |
| --- | --- | --- | --- | --- | --- | --- | --- |
| virus | sample days | S205 | N205 | K178 | R178 | R649 | K649 |
| WI.97.ic | 8, 8, 10, 10 | 100% (4/4) | 0% (0/4) | 100% (4/4) | 0% (0/4) | 100% (4/4) | 0% (0/4) |
| NY.19.32 | 5, 6, 10, 10 | 0% (0/4) | 100% (4/4) | 0% (0/4) | 100% (4/4) | 0% (0/4) | 100% (4/4) |
| NY.19.12 | 6, 7, 10, 10 | 0% (0/4) | 100% (4/4) | 0% (0/4) | 100% (4/4) | 0% (0/4) | 100% (4/4) |

Thus, morbidity, mortality, and tissue tropism within the brain and spleen at six days post-infection and endpoint were assessed for WI.97.ic^NY.19.12^ compared to NY.19.12 and WI.97.ic. Consistent with previous experiments, there was minimal weight loss by day six post-infection, but by day 12 post-infection, mice in all cohorts had either lost weight and met criteria for euthanasia or recovered (**Figure 8A**). There were no significant differences in survival between strains (**Figure 8B**), but at day four post-infection, ∼70% of WI.97.ic^NY.19.12^ and NY.19.12 infected mice displayed mild clinical signs compared to less than 20% of WI.97.ic infected mice (**Figure 8C**). By day six post-infection, there were higher rates of neurologic clinical signs in NY.19.12 and WI.97.ic^NY.19.12^ infected mice compared to WI.97.ic infected mice (**Figure 8C**). Additionally, there was significantly more forelimb weakness in NY.19.12 and WI.97.ic^NY.19.12^ cohorts compared to WI.97.ic infected mice (p<0.05) (**Figure 8D**).

**Figure 8.**
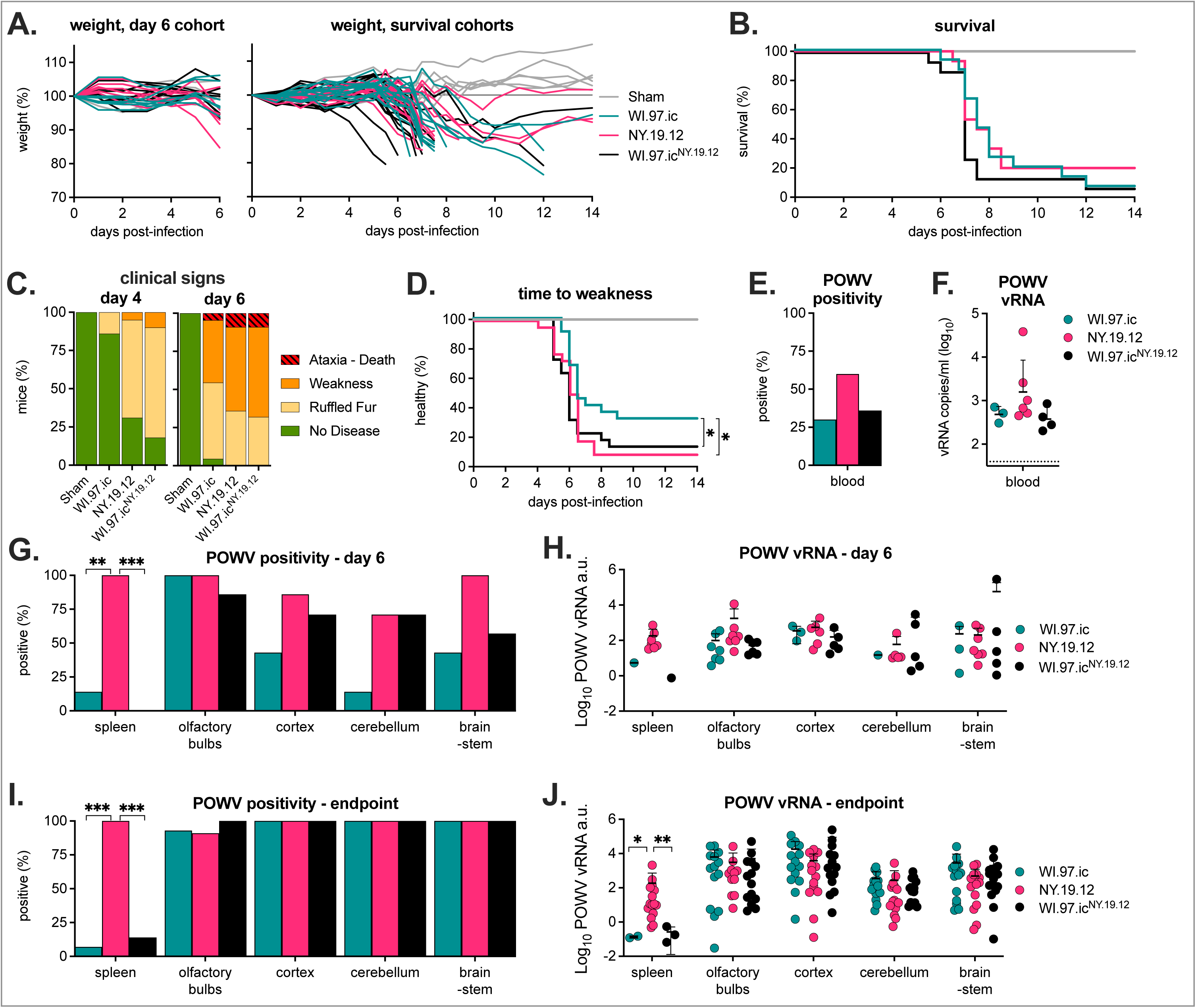
WI.97.ic^NY.19.12^ infected mice display varying trends in disease progression and virus distribution in the brain at six days post-infection and at endpoint. Mice were S.C. inoculated with 10^3^ PFU POWV (WI.97.ic^NY.19.12^, NY.19.12, and WI.97.ic) or sham and sacrificed at either day six post-infection (n = 7 per virus) or at endpoint (n = 15 total, two independent experiments, n = 8 and n = 7 per virus). **A)** Percent weight change from starting weight for day six cohort and endpoint cohorts. **B)** Kaplan-Meier survival curves for endpoint cohorts, no significant differences between viruses with log-rank (Mantel-Cox) test (p>0.05)**. C)** Clinical signs four and six days post-infection for all cohorts combined (n = 22 per virus). **D)** The amount of time for mice to exhibit forelimb weakness. Viruses were compared using log-rank (Mantel-Cox) test, * p<0.05. **E)** POWV positive (%) blood samples and **F)** vRNA levels from day two post-infection (day six cohort, one endpoint cohort, n = 14 per virus). The dotted line shows the limit of detection. Positive POWV (%) of **G)** day six and **I)** endpoint cohort spleen, olfactory bulbs, cortex, cerebellum, and brainstem (Fisher’s exact test, ** p<0.01, *** p<0.001). Log_10_ POWV vRNA normalized to GAPDH intracellular levels from spleen, olfactory bulbs, cortex, cerebellum, and brainstem for **H)** day six cohort and **J)** endpoint cohorts. a.u. = arbitrary units. Within each sample type, two-way ANOVA with Tukey’s multiple comparisons test (* p<0.05, ** p<0.01).

Higher rates of viremia and higher vRNA levels in the blood were observed at two days post-infection in NY.19.12 infected mice compared to both WI.97.ic and WI.97.ic^NY.19.12^ (**Figure 8E/F).** By day six post-infection, vRNA was detected at comparable rates in the olfactory bulbs across strains. Interestingly, WI.97.ic^NY.19.12^ displayed an intermediate infection phenotype between NY.19.12 and WI.97.ic, with higher rates of infection in the cortex, cerebellum, and brainstem compared to WI.97.ic (**Figure 8G**). Levels of vRNA within each brain tissue were comparable across strains (**Figure 8H**). By endpoint, vRNA was detected at similar frequencies and levels in the olfactory bulbs, cortex, cerebellum, and brainstem samples across infection cohorts (**Figure 8I/J**). Notably, compared to both WI.97.ic and WI.97.ic^NY.19.12^, vRNA was detected at significantly higher rates in the spleen of NY.19.12 infected mice at both day six post-infection and endpoint, and had significantly higher levels of vRNA by disease endpoint (**Figure 8G/I/J**).

Infected mice brains were then processed for IFA and stained for POWV antigen (**Figure 9A-C)**. Notably, NY.19.12 and WI.97.ic^NY.19.12^ were both detected in the olfactory bulbs, cerebellum, and brainstem. However, WI.97.ic^NY.19.12^ and WI.97.ic had POWV infection in the hypothalamus. All infection groups shared POWV distribution patterns in the olfactory peduncle.

**Figure 9.**
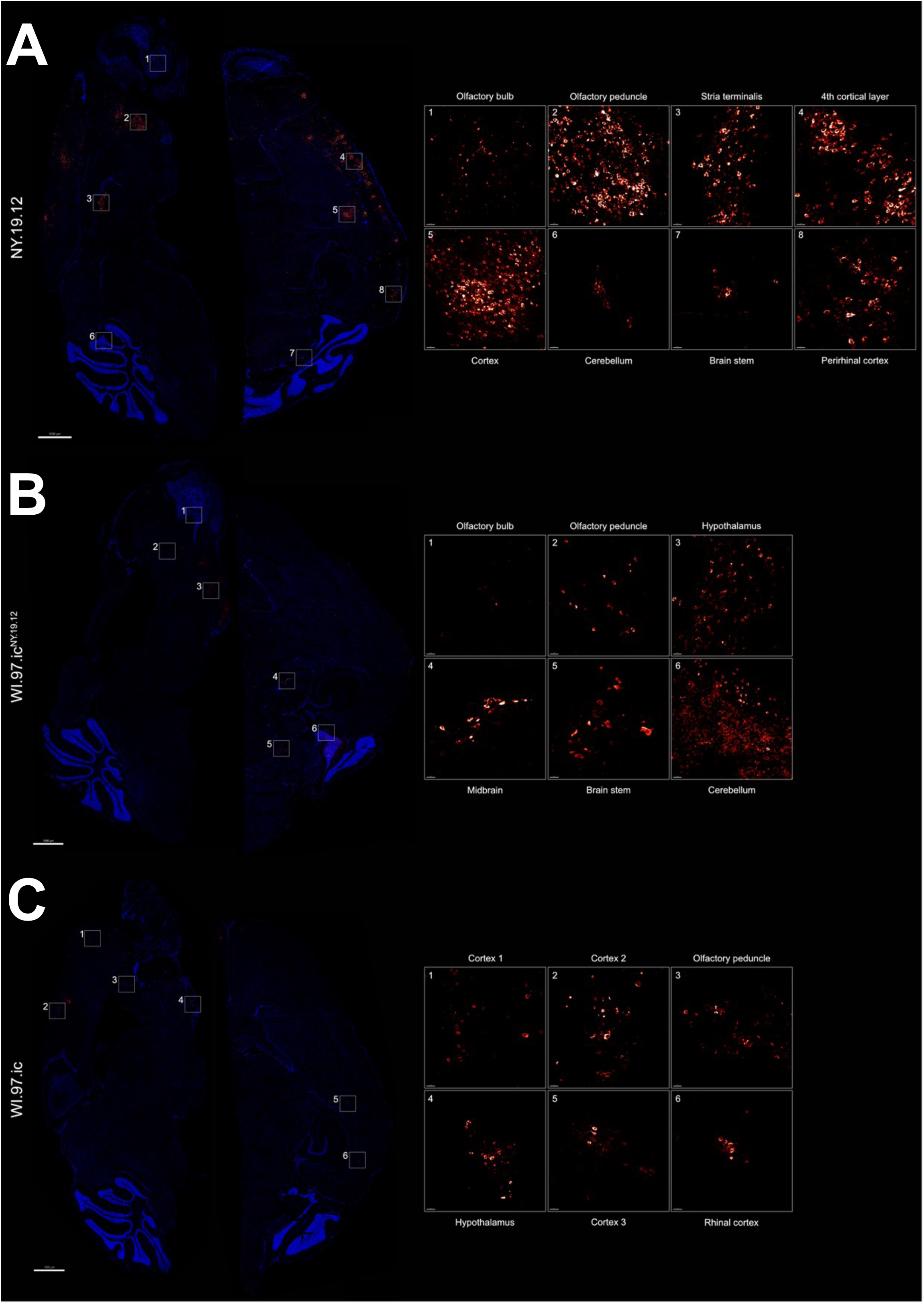
POWV viral distribution in the brain of NY.19.12, WI.97.ic^NY.19.12^, and WI.97.ic infected mice. Mice were S.C. inoculated with POWV (WI.97.ic^NY.19.12^, NY.19.12, and WI.97.ic) or sham and sacrificed at day six post-infection (n = 7 per virus). Brains were processed for IFA and stained for POWV in red. Representative sagittal images are shown. Zoomed in boxes show detection of POWV antigen within specific brain regions. **A)** NY.19.12 infected mouse brain with POWV infection in the 1) olfactory bulb, 2) olfactory peduncle, 3) stria terminalis, 4) 4^th^ cortical layer, 5) cortex, 6) cerebellum, 7) brain stem, and 8) perirhinal cortex. **B)** WI.97.ic^NY.19.12^ infected mouse brain with POWV infection in the 1) olfactory bulb, 2) olfactory peduncle, 3) hypothalamus, 4) midbrain, 5) brain stem, and 6) cerebellum. **C)** WI.97.ic infected mouse brain with POWV infection in the 1-2) cortex, 3) olfactory peduncle, 4) hypothalamus, 5) cortex, and 6) rhinal cortex.

## Discussion

We investigated several POWV lineage II strains from New York, Massachusetts, and Wisconsin to define the extent of virus strain-dependent variation in POWV pathogenesis. To capture the diversity of POWV lineage II circulating in the United States, we selected strains that span the geographic range of reported human cases. Each strain was originally isolated from ticks and maintained with minimal *in vitro* passaging. Genomic sequencing confirmed that these isolates are representative of viruses currently enzootic in their respective regions. We also used rescued virus from a well-characterized infectious clone of the ‘Spooner’ strain as a lineage II reference virus (WI.97.ic).^14^ Taken together, the panel of viruses used here provides a robust and relevant sample of the genetic and spatiotemporal diversity of POWV lineage II.

Variation in pathogenesis within lineage I strains, and differences between lineage I and II, have been documented.^13^ However, the extent of variation in pathogenesis within lineage II is unknown. Viral genomics studies have demonstrated extensive genetic diversity within POWV lineage II, but most studies of POWV lineage II use the Spooner isolate with even fewer studies using other lineage II isolates.^15–18^ Our studies revealed significant variation in morbidity and mortality in mice, with some common features of pathogenesis that are consistent across all POWV lineage II strains. This data suggests that, regardless of strain, 1) C57BL/6 succumb to disease within 7-14 days 2) weight loss occurs upon neurological disease onset, and 3) by disease endpoint, there are high levels of vRNA throughout the brain. In addition, histopathology of NY.19.12 and WI.97.ic infected mice that succumbed to disease showed similar severity and distribution of inflammation in the brain. These observations align with previous research on POWV infection in mice, which typically results in severe neurological disease within 14 days of infection.^19^ However, rates of survival vary greatly by strain after S.C. inoculation. Interestingly, NY.19.12 and NY.19.32 caused more severe disease (weight loss, clinical signs) and earlier vRNA detection in the serum and brain. These results show important similarities within lineage II as well as differences in rates of tissue infection and virus distribution in the brain at early timepoints.

Genetic analysis revealed three unique amino acid mutations shared by NY.19.12 and NY.19.32 in the E, NS1, and NS5 proteins. These mutations were also found in a subset of lineage II strains that cluster phylogenetically, indicating they likely provide some fitness advantage causing them to be maintained in nature. Although genetically restricted infectious clone-derived viruses are often attenuated relative to naturally isolated strains composed of genetically diverse quasispecies^20,21^, they are an important tool allowing direct comparisons between mutants. In this study, WI.97.ic^NY.19.12^ exhibited comparable tissue tropism to the natural NY.19.12 strain, with similar vRNA levels in the cerebellum and similar distribution in several regions of the brain, suggesting that these amino acid mutations contribute to neuroinvasion. However, the mechanisms underlying neurotropism and how the host response shapes viral distribution within the brain are not well defined. These results showed that NY.19.12 primarily targets neurons with limited astrocyte recruitment, which is consistent with current POWV literature and contributes to the growing understanding of POWV infection within the CNS.^10,13,22^ Although there have been some studies characterizing POWV neuropathology and immune involvement in the CNS^22,23^, the field remains relatively understudied. Therefore, further high-resolution imaging of WI.97.ic^NY.19.12^ in the brain is warranted to assess neuronal infection and astrocyte recruitment in comparison to NY.19.12.^11,19^ To better define regional neuroinvasion and pathogenesis, recent studies have characterized other tick-borne flaviviruses using whole-brain imaging approaches. For example, using this approach, the Överby group demonstrated that both the E, NS1, and NS5 proteins may contribute to tick-borne encephalitis virus neurotropism. Such techniques challenge existing methods and literature, ultimately allowing for a deeper understanding of cell-type specific responses and viral tropism for tick-borne flaviviruses such as POWV.^11,12^

Some disease phenotypes, such as viral persistence in the spleen, were not recapitulated by WI.97.ic^NY.19.12^, suggesting that additional viral factors may influence pathogenesis outside of the mutations engineered into the infectious clone. Although spleen tropism in flaviviruses is not well understood, it was found that POWV lineage I infected mice had prominent infection of splenic macrophages.^10^ Similarly, studies on dengue virus have demonstrated the spleen can serve as a site for viral replication and contribute to disease severity.^24,25^ Moreover, our group has shown that some Northeast and Midwest lineage II isolates have 100% infection of the spleen throughout early time points.^15^ This indicates that enhanced spleen infection may not be unique to NY.19 strains. One explanation could be the absence of quasispecies diversity of infectious clone-derived virus, although how this impacts spleen tropism is unclear. However, studies among tick-borne flaviviruses have shown that infectious clones often have reduced neurovirulence because they lack population diversity, and this could be an explanation for the phenotypic differences in the spleen as well.^21,26^ Further studies are required to further understand the importance of the spleen in POWV infection.

Additionally, other factors may contribute to phenotypic differences between WI.97.ic^NY.19.12^ and NY.19.12. These include synonymous mutations that affect RNA structure, undescribed determinants in the 3’ and 5’ untranslated regions of the genome, or other single nucleotide polymorphisms (SNPs).^27–29^ While the impact of POWV SNPs on disease phenotypes remains largely unknown, recent studies have begun to elucidate the role of SNPs in flavivirus pathogenesis. In Japanese encephalitis virus, a single amino acid change in E was linked to increased neurovirulence^29^, and research on tick-borne encephalitis virus found four amino acids in NS5 led to increased neurovirulence.^30^ Although these SNPs are not the same amino acid mutations in this study, it is important to note that SNPs can be critical for flavivirus disease phenotypes. For many flaviviruses, glycosylation mutations in the E protein can impact virulence and pathogenesis; although E-205 is on the surface of E, it is not associated with any glycosylation sites.^31^ These studies suggest there are likely multiple genetic determinants that contribute to POWV disease and neuroinvasion. Importantly, non-coding genetic variants, such as those affecting regulatory regions, microRNAs, or splicing elements, may also play a critical role in modulating host susceptibility and viral pathogenesis.^32^ Future studies incorporating transcriptomic and epigenomic analyses could help clarify these contributions. While factors dictating POWV neuroinvasion remain largely undefined, these results indicate that SNPs or non-coding RNAs may contribute to shaping variation in viral pathogenicity. Overall, this study provides valuable insights into the factors influencing POWV lineage II disease progression and tissue tropism.

## Materials and Methods

### Cells and viruses

BHK-21 (ATCC CCL-10) and HEK293T (ATCC CRL-3216) cells were grown in Dulbecco’s modified eagle medium (DMEM) supplemented with 10% fetal bovine serum (FBS), 10 units/mL penicillin, 10 μg/mL streptomycin at 37°C and 5% CO_2_. Viruses (described in **Table 1**) were passaged on BHK cells, supernatant harvested 3-6 days post-infection, and frozen at -80°C prior to use. Virus inoculum for mouse studies was backtitered via plaque assays to validate virus concentration.

### Plaque assays

Standard plaque assays were used to determine infectious virus concentrations. BHK cells were seeded one day prior to infection to be confluent the following day. Virus samples were thawed, serially diluted in infection media (standard growth media with 2% FBS), added to cells and incubated at room temperature for 1 hour on a plate rocker. Cells were overlaid with semi-solid 0.6% tragacanth media and incubated at 37°C for five days. Cells were fixed and stained with 20% ethanol (EtOH) and 0.1% crystal violet. Plaques were counted manually to calculate titer.

### POWV mouse infections

C57BL/6 mice (strain #000664) were obtained from Jackson Laboratories. Approval for animal protocols was obtained by the Colorado State University Institutional Animal Care and Use Committee (protocol #5074). All animal infections were conducted in Colorado State University ABSL-3 containment. 8-week-old female mice were S.C. inoculated with 10^3^ PFU POWV in 100uL or sham infected with 100uL DMEM. Mice were weighed and clinical signs were recorded unblinded by the same individual throughout the course of the studies. Drastic weight loss was observed in the first mouse study (see **Figure 1**) due to food access – the pellets in the feeder are difficult to reach for the mice upon clinical sign onset (ruffled fur, hunched posture). For subsequent studies, food pellets were placed in the bottom of the cage once clinical sign onset was observed. Clinical sign criteria were categorized as listed in **Table 3**. Euthanasia criteria were met when clinical signs of ataxia were observed or at 20% weight loss.

**Table 3.** Clinical sign criteria for POWV infected mice.

| Clinical signs |
| --- |
| Healthy |
| Hunched posture, ruffled fur |
| Forelimb weakness |
| Ataxia |

### Mouse necropsies and sample processing

Submandibular blood collection was performed with 4 mm lancets and blood was stored in BD Microtainer tubes with serum separator additive (serum) or EDTA (whole blood). For necropsies, mice were euthanized via 5% isoflurane and cervical dislocation, and blood was collected via cardiac puncture. Cortex and spleens were harvested. For brain region analysis using immunofluorescence, mice were deeply anesthetized with 5% isoflurane and perfused with 30 mL of PBS then 30 mL of freshly prepared 4% paraformaldehyde, and brain regions were harvested. For brain region analysis via qRT-PCR, mice were deeply anesthetized with 5% isoflurane and perfused with 30 mL of PBS and brain regions were harvested. In the time course study (see **Figure 3**), samples were placed into 2 mL snap cap tubes with a metal bead, and 10% of each tissue weight was added in volume with DMEM and frozen at -80°C until homogenization. For subsequent studies, samples were placed into 2 mL snap cap tubes with 1 mL of 1X DMEM and a metal bead and frozen at -80°C until homogenization. Mouse tissues were rapidly thawed and homogenized at 30 Hz/s for 2 minutes, then centrifuged at maximum speed for 10 minutes to pellet cellular debris.

### Nucleic acid extraction and qRT-PCR

Viral nucleic acid was extracted from 50 μL of serum, whole blood or homogenized mouse tissue using the Omega Viral DNA/RNA Extraction Kit on the KingFisher Flex instrument following manufacturer’s instructions. Samples were either normalized for qRT-PCR via 10% weight/volume per sample before homogenization (see **Figure 3**) or to total RNA concentration after homogenization (see **Figure 5, 8**). Quantitative real-time PCR (qRT-PCR) was performed using the EXPRESS One-Step qRT-PCR kit and POWV primer-probes targeting the NS5 coding region (blood, spleen, olfactory bulbs, cortex, cerebellum, brainstem), or EXPRESS One-Step SYBR GreenER Kit with mouse GAPDH primers (spleen, olfactory bulbs, cortex, cerebellum, brainstem), in duplicate (**Table 4**).^33^ vRNA copies were determined using a POWV RNA standard covering the NS5 gene as previously described (**Figure 3** and blood/serum samples) or normalized using the delta delta CT method (normalized to GAPDH) (**Figure 5, 8**).^15^ Details provided in figure legends.

**Table 4.**
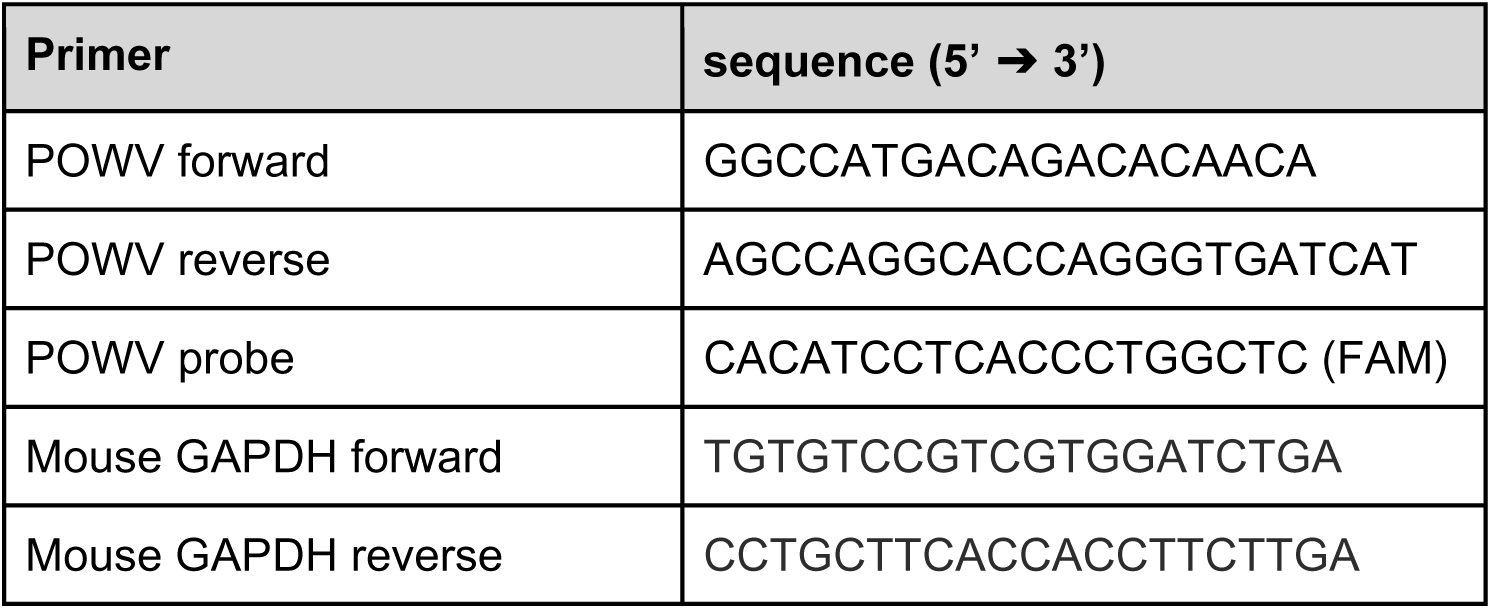
Primer-probe sequences for qRT-PCR assays.

### Sanger sequencing of mutations

vRNA was extracted as described above, and reverse transcription and PCR performed using QIAGEN OneStep RT-PCR Kit following manufacturer’s protocol and primers to amplify regions containing each of the mutations (E, NS1 and NS5) (**Table 5**). PCR products were gel purified and cleaned up using MACHERRY-NAGEL NucleoSpin Gel and PCR Clean-up kit, then Sanger sequenced.

**Table 5.**
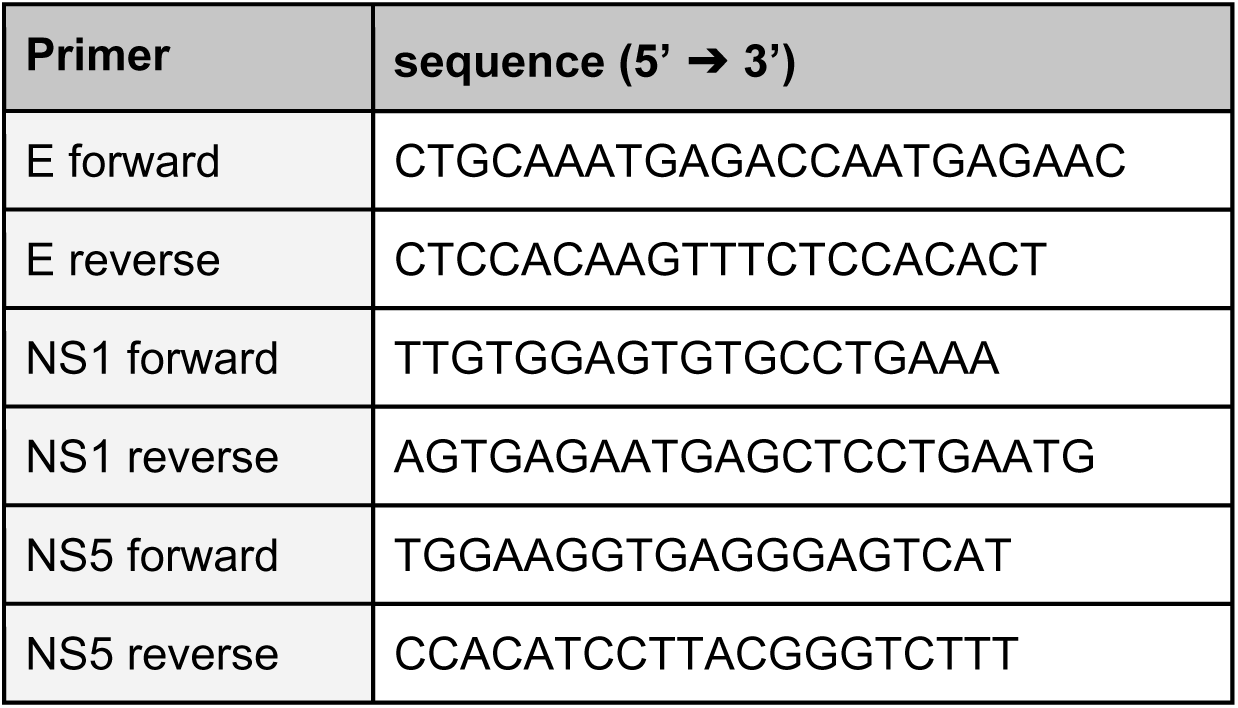
Primer sequences for Sanger sequencing.

### Generation of mutant infectious clone

The POWV lineage II two-plasmid cDNA infectious clone (WI.97.ic) was obtained from Aaron Brault, CDC and was rescued and propagated as previously described.^14^ E and NS1 mutations were engineered into plasmid 1 via NEBuilder HiFi DNA Assembly Cloning Kit, and the NS5 mutation was engineered into plasmid 2 via NEB Q5® Site-Directed Mutagenesis Kit, using primers designed via NEBuilder Assembly Tool. Mutant infectious clone (WI.97.ic^NY.19.12^) plasmids were transformed into Stbl3 *E. coli* and grown on ampicillin-treated agar plates at 37°C overnight. Individual colonies were isolated and grown overnight in liquid culture and isolated and purified via QIAprep Spin Miniprep Kit. Whole plasmids were sequence verified using Plasmidsaurus. Plasmids 1 and 2 were linearized with BstZ171 and ligated together at a 1:3 ratio with T4 DNA ligase. The ligated product was linearized with NotI, and RNA transcribed via mMESSAGE mMACHINE™ T7 Transcription Kit. Full-length WI.97.ic^NY.19.12^ RNA was transfected into HEK293T cells using Lipofectamine 3000. Supernatant was collected at approximately 4-5 days post transfection once cells reached ∼50% cytopathic effect. Viral stocks were passaged on BHK-21 cells, concentrated via centrifugation, and titered by plaque assay. Virus sequence at each mutation site (E, NS1, and NS5) was confirmed via Sanger sequencing as described above.

### Histopathology

Mouse heads were fixed in 10% paraformaldehyde and then decalcified for 48 hours in nitric acid. Sagittal sections of the skull were then paraffin embedded, routinely processed, and stained with hematoxylin and eosin (H&E) for histologic evaluation.

### Immunohistochemistry

The right hemisphere from brains of infected mice were dehydrated in 30% sucrose frozen in OCT Embedding matrix and sectioned (sagittal and axial sections) at 10 μm thickness using CryoStar NX50 cryostat according to manufacturer’s instructions at -20°C. Slides were thawed at 37°C for 5 minutes, hydrated in PBS followed by permeabilization and blocking using PBS, 0.2% TritonX, 10% goat serum and 0.1% NaN3. Sections were stained with primary antibodies against POWV E protein (GeneTex, GTX640340, 1:1000), POWV NS3 protein (GeneTex, GTX642219, 1:1000), astrocyte marker GFAP (Abcam, ab4674, 1:500) and secondary antibodies donkey anti-rabbit A488 (Invitrogen, A21206, 1:500), donkey anti-rabbit A647 (Invitrogen, A31573, 1:500), DAPI (Sigma, 1:1000) and neuronal marker NeuN-A647 (Abcam ab190565, 1:200). Slides were mounted and imaged by Leica SP8 confocal microscope using a HC PL APO 20x/0.75 Dry objective. Images were created using Imaris software (version 10.1.1, Bitplane, Zürich, Switzerland).

## Data analysis and statistics

All data were analyzed using GraphPad Prism Version 9.5.0. Statistical tests are described in figure legends. Sequences were analyzed in Geneious Prime ® 2023.0.1.

## Data availability

All data are available upon request.

## Acknowledgements

We thank Aaron Brault (CDC) for providing the POWV lineage II infectious clone. This research was funded by the National Institute of Allergy and Infectious Diseases (R01AI137424) (G.D.E.), the Laboratory for Molecular Infection Medicine Sweden (MIMS) (A.K.Ö.), Umeå Center for Microbial Research Visiting Scientist program (A.K.Ö.) Umeå University Medical Faculty Research Sabbatical (A.K.Ö.), the Swedish Research Council grant (2020-06224) (A.K.Ö.). SC was supported by funding from National Institute of Allergy and Infectious Diseases (T32AI162691). We thank CSU Laboratory Animal Resources staff and facilities for support (RRID: SCR_022157). We also acknowledge the Biochemical Imaging Center at Umeå University (BICU) and the National Microscopy Infrastructure for microscopy support (NMI; VR-RFI 2019-00217). Figures were generated using BioRender.

## Author contributions

SJC: Conceptualization, funding acquisition, investigation, writing - original draft. ENG: Investigation, writing - review and editing. EN: Investigation. CET: Investigation. KXK: Investigation. ACF: Investigation. AV: Investigation. AKÖ: Funding acquisition, investigation, methodology, supervision. GDE: Funding acquisition, writing - review and editing, supervision. All authors revised the final manuscript.

